# CBASS limits bacteriophage production while maintaining cell viability in *Pseudomonas aeruginosa*

**DOI:** 10.64898/2026.02.24.707611

**Authors:** Erin Huiting, Bruce Wang, Esther Shmidov, Sriharshita Musunuri, Zemer Gitai, Joseph Bondy-Denomy

## Abstract

CBASS is an immune pathway that recognizes phage infection and generates cyclic nucleotide signals, which directly activate effectors that stop phage replication. Membrane-acting effectors are proposed to induce cell death to prevent phage replication; however, this mechanism has not been assessed with endogenous expression levels of the effector. We therefore sought to determine the cell viability outcomes of the CBASS phospholipase effector (CapV) upon activation with 3’,3’-cGAMP signals in *Pseudomonas aeruginosa*. Here, we surprisingly observe that constitutive 3’,3’-cGAMP signaling from the synthase (CdnA) enables robust cell growth and viability while effectively abolishing phage production in a CapV-dependent manner. Exogenous 3’,3’-cGAMP also enhances CBASS anti-phage activity and cell growth. Moreover, constitutive activation of the CapV effector induces no cell fitness cost, and blocks replication of many, but not all, phages. This demonstrates that a cyclic nucleotide-activated CBASS effector possess a degree of phage specificity that has been previously overlooked. When CBASS is active, phage transcription and initiation of DNA replication proceed normally, but phages do not reach maximum DNA levels and fewer mature virions are produced. Based on these findings, we propose that CapV interferes with the early stages of phage capsid assembly at the cell membrane and resultantly disrupts DNA packaging. Collectively, we demonstrate that a successful CBASS response antagonizes a late-stage of the phage replication cycle while maintaining cell viability.

## Introduction

Cyclic GMP-AMP synthases (cGAS) are an evolutionarily conserved enzyme family, with members that can produce one or more unique cyclic oligonucleotides. Thousands of cGAS-like homologs have been identified in animals^1–3^ and bacteria^4–8^ that produce at least 16 signals. These bacterial cGAS/DncV-like nucleotidyltransferases (CD-NTases) produce signals that activate distinct effector proteins, which limit phage replication and protect the surrounding bacterial cells^9–14^, collectively referred to as cyclic-oligonucleotide-based anti-phage signaling systems (CBASS). To protect themselves, phages have evolved mechanisms to overcome CBASS^15–24^. In response, the bacterial host may increase or modify CD-NTase activity to overcome these phage inhibitory mechanisms^17,25^. Currently, it is proposed that CBASS effectors induce cell death or dormancy, but all anti-phage studies heterologously express diverse CBASS operons within phage-sensitive *Escherichia coli* strains that naturally lack CBASS^8,10,11,13,14,26^. However, the cell viability outcome of CBASS during phage infection has yet to be studied under endogenous expression levels of the effector.

The first active CD-NTase (known as DncV) was studied in *Vibrio cholerae*, where 3’,3’-cGAMP signals activate a phospholipase effector (VcCapV) that degrades phospholipids *in vitro*^27^. Activation was achieved through DncV overexpression, and in later studies through folate depletion, both of which induced a cell death phenotype^27,28,34–36^. Most recently, the combination of DncV overexpression with *Acinetobacter baumannii* CapV effector (AbCapV), also induces a cell death phenotype *in vivo* and AbCapV cleavages of lipid substrates *in vitro*^25^. In our previous study, we identified a strain of *Pseudomonas aeruginosa* featuring an anti-phage CD-NTase and CapV effector that is homologous to the DncV system in *V. cholerae*. The *P. aeruginosa* CD-NTase (CdnA) is a part of Type II-A CBASS and produces 3’,3’-cGAMP, which activates the corresponding CapV (PaCapV) effector and reduces the titer of phage PaMx41 over 10,000-fold^16^. PaCapV also directly binds to 3’3’-cGAMP, which activates the cleavage of lipase substrates *in vitro*^16^. Here, we used this same *P. aeruginosa* strain (Pa011) to determine how the activated PaCapV effector prevents phage replication. Using strains with constitutively high 3’,3’-cGAMP signaling, through an additional copy of CdnA or exogenous 3’,3’-cGAMP, we observed that activated CapV limits phage replication and cells grow better at all phage infection levels compared to cells lacking CBASS. These findings demonstrate that activated CapV does not induce a cell fitness cost. We also identify classes of phage that are sensitive or resistant to the activated CapV effector, introducing the concept of CBASS effector specificity. For CBASS-sensitive phage, maximum DNA levels are not reached, and phage virion production is reduced 10-100-fold within a single round of infection. As an alternative model to CapV-induced cell death, we propose that CapV interferes with early stages of capsid assembly (i.e. the procapsid) at the cell membrane and disrupts subsequent phage DNA packaging. Our collective findings demonstrate that CBASS activity limits phage replication while maintaining cell viability, which presents a significant shift in our current understanding of CBASS membrane effectors and their cellular outcomes.

## Results

### Increasing CdnA activity potently targets phage

Type II-A CBASS in *Pseudomonas aeruginosa* strain BWHPSA011 (Pa011) limits the replication of dsDNA lytic phage PaMx41 lacking the CBASS inhibitor Acb2, but PaMx41 expressing Acb2 can overcome CBASS activity (Figure 1A-B)^16^. It was independently proposed that increasing 3’,3’-cGAMP production can overwhelm CBASS inhibitors^17^. We initially sought to test this model, but inadvertently generated an ideal strain to examine the mechanism of the CapV effector. To increase 3’3’-cGAMP production, we overexpressed CdnA in the Pa011 strain under an inducible promoter, which generated a constitutively active CdnA enzyme with endogenous CapV levels (Figure 1A-B). This CdnA overexpression strain targeted both PaMx41Δ*acb2*, and an isogenic PaMx41 phage expressing *acb2*, reducing their titers an additional two or three orders of magnitude in a CapV-dependent manner (Figure 1C, S1A). CdnA overexpression in Pa011 resulted in ∼6.5 µM of intracellular 3’,3’-cGAMP, which was detected in the presence or absence of phage (Figure 1D), demonstrating constitutive CBASS activity. In contrast, in the Pa011 strain with endogenous CdnA levels, ∼0.013 µM of 3’,3’-cGAMP was detected only in the presence of phage infection, but no 3’,3’-cGAMP was detected in the absence of phage (Figure 1E). Therefore, these data show that constitutive CBASS activity more potently stops phage infection through enhanced 3’,3’-cGAMP signal production, which we used next to observe the cellular outcome of CapV-dependent activity.

**Figure 1.**
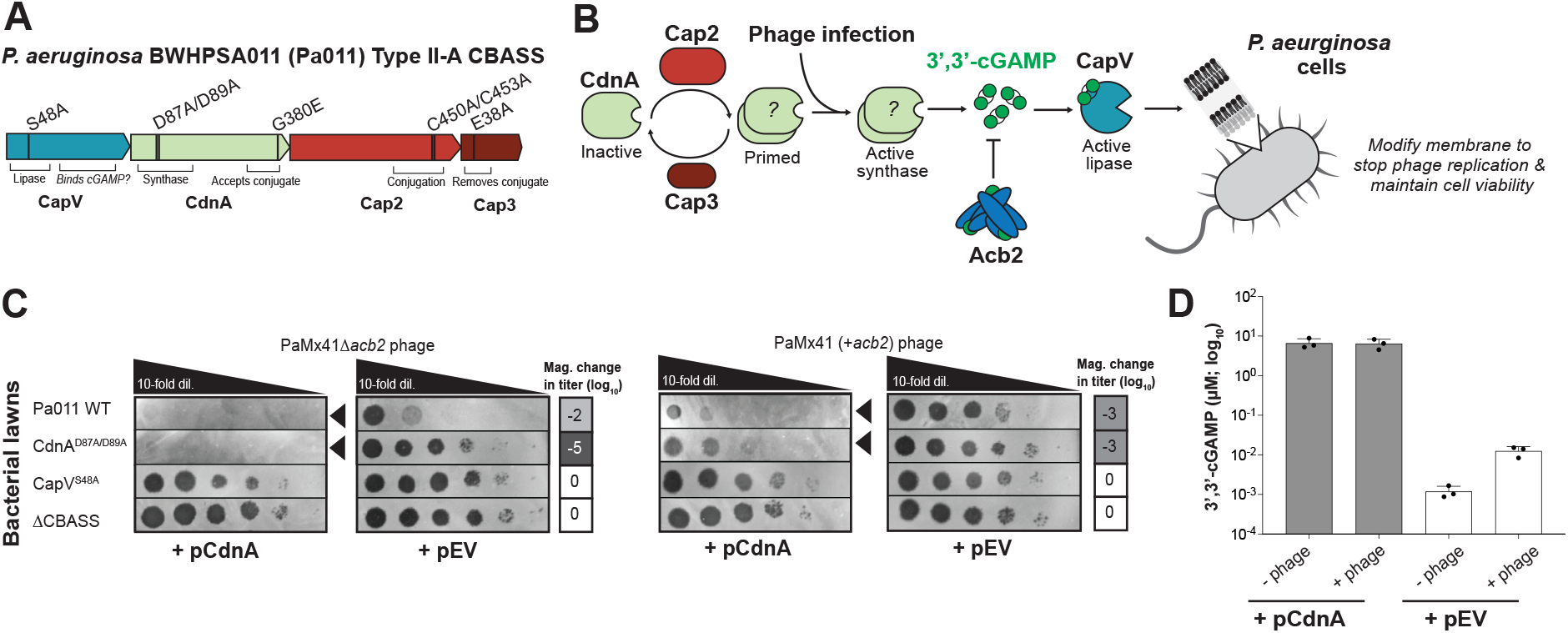
Increasing CdnA activity potently targets PaMx41Δacb2 and +acb2 phage. **(A)** Pseudomonas aeruginosa strain BWHPSA011 (Pa011) Type II-A CBASS operon with domains and functional mutants indicated. **(B)** Working model of Pa011 Type II-CBASS, wherein CdnA produces a 3’,3’-cGAMP signals, which the Acb2 inhibitor may sequester, or the signals directly bind to and activate the CapV effector. We then propose that the CapV effector modifies the cell membrane to disrupt phage assembly and effectively stopping phage replication while maintaining cell viability. **(C)** PaMx41 phage that is CBASS sensitive (Δacb2) or CBASS resistant (+acb2) spotted across lawns of Pa011 WT, CdnA or CapV catalytically dead, and ΔCBASS strains expressing CdnA or empty vector (EV) plasmids (0.1% arabinose, n=3). These assays were used to quantify the order of magnitude change in phage titer by comparing the number of spots (with plaques, or clearings if plaques were not visible) on the CdnA expressing strain divided by the EV strain. Black arrowheads highlight significant reductions in phage titer. **(D)** Quantification of intracellular 3’,3’-cGAMP in Pa011 CapVS48A strains expressing CdnA or EV plasmids within the absence or presence of PaMx41Δacb2 phage at 60 minutes post-infection (MOI: 5, 0.1% arabinose; n=3).

### Type II-A CBASS maintains cell viability upon phage infection

To assess the cellular viability of CBASS anti-phage activity, we initially used a standard liquid infection assay to measure bacterial population growth over the course of many hours. During infection of the Pa011 strain with endogenous CBASS activity (Pa011 WT), we observed a partial decrease in cell density as the multiplicity of infection (MOI) of PaMx41Δacb2 increased from 0.2 to 20 (Figure 2A, S2A). While the reduced growth may have been due to an abortive infection mechanism, we instead observed partial phage replication in this Pa011 WT strain. Over the course of eight hours, phage virion production was reduced 10-100-fold compared to the strain lacking CBASS, but virion production was not fully abolished (Figure 2B). To assess an abortive cell death outcome, it’s common to use a rapid decrease in OD600 values, but the partial failure of CBASS immunity at high MOI limited our ability to examine the cellular outcome of CBASS activity. We therefore leveraged the Pa011 strain with constitutive CBASS activity (e.g. CdnA overexpression, but endogenous CapV levels) and its robust anti-phage activity to assess whether a stronger CBASS immunity manifests as abortive. To this end, we hypothesized that constitutively high CBASS activity would enhance an abortive cell death outcome. Surprisingly, the Pa011 cells with constitutively active CBASS grew remarkably well in the presence of low and high multiplicities of PaMx41Δ*acb2* phage infection similarly to uninfected cells (MOI 0.2-20; Figure 2A). These collective data suggest that activated CapV does not induce a cell death response to stop phage replication.

**Figure 2.**
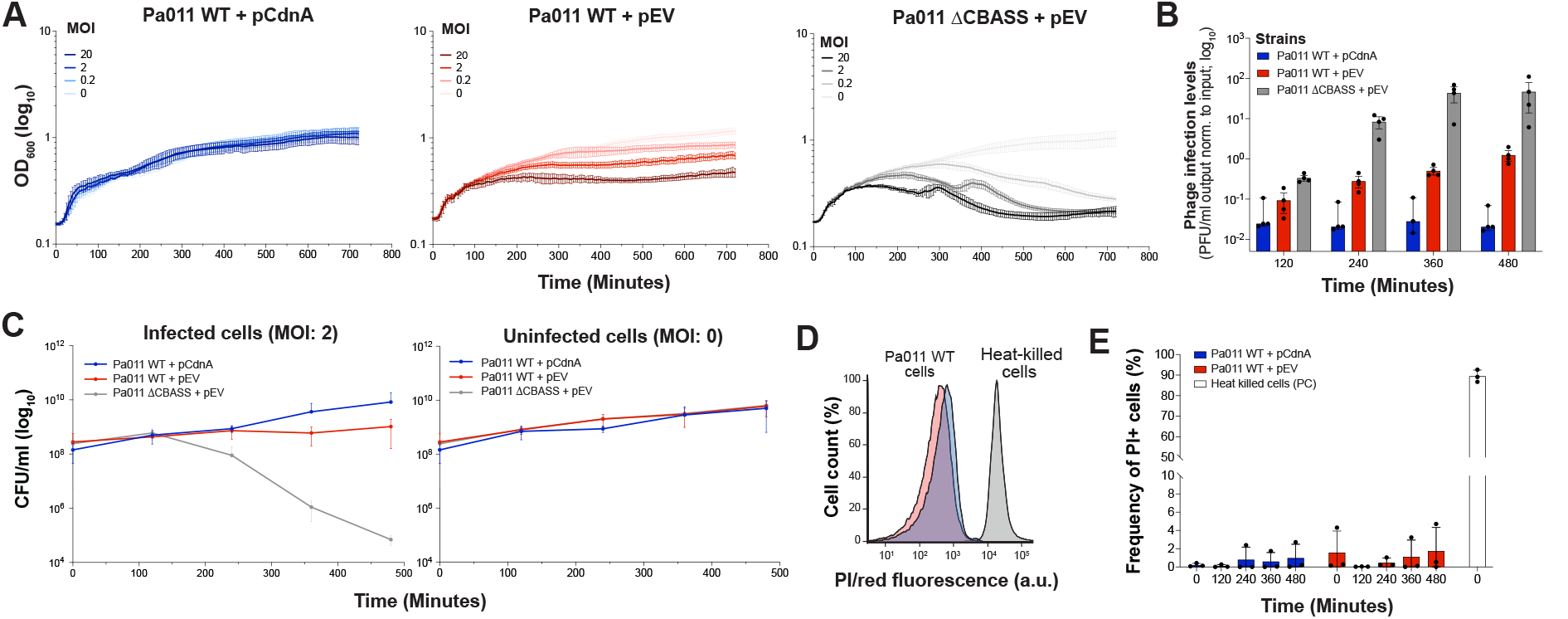
Type II-A CBASS maintains cell viability during PaMx41Δacb2 phage infection. **(A)** OD600 measurements of Pa011 strains expressing CdnA or empty vector (EV) plasmids infected with an increasing multiplicity of infection (MOI) of PaMx41Δacb2 phage over 720 minutes (0.05% arabinose, n=3 +/-s.d.). **(B)** Phage infection levels (PFU/ml of output normalized to input phage) of Pa011 WT and ΔCBASS strains expressing CdnA or EV plasmids infected with PaMx41Δacb2 phage over 720 minutes (MOI: 2, 0.05% arabinose, n=3 +/-s.d.). **(C)** Colony forming units (CFU/ml) of infected and uninfected Pa011 strains. **(D)** Histogram representation of Propidium Iodide (PI) negative (live) or positive (dead or porous) cells of infected Pa011 strains at 480 minutes post329 infection. Heat-killed cells as used as a positive control (PC) because they effectively retain PI dye. **(E)** Frequency of PI positive cells over 480 minutes of phage infection. Assays performed in D-E used PaMx41Δacb2 phage (MOI:2, 0.05% arabinose; n=3 +/-s.d.)

To determine whether increased cell growth is due to viable cells effectively stopping phage replication, we initially measured bacterial cell viability and phage titer over several hours using an intermediate MOI of 2 to ensure that most bacterial cells would be infected. In Pa011 strains with endogenous or constitutive CBASS activity, the bacterial cell count remained stable at ∼10^8^ CFU/ml and then increased to ∼10^9^ CFU/ml or ∼10^10^ at eight hours post-infection (Figure 2C). While the starting bacterial cell count was the same for the Pa011 ΔCBASS strain, the cell counts dropped starting at four hours post-infection and then stabilized at ∼10^5^ CFU/ml (Figure 2C), demonstrating the PaMx41 phage can slowly, albeit effectively, lyse >99.9% of cells in the absence of CBASS. Furthermore, the Pa011 strains with functional CBASS reduced phage titer one to four orders of magnitude over this time course in comparison to the Pa011 ΔCBASS strain (Figure 2B), validating that CBASS anti-phage immunity is active.

To further study the viability of single bacterial cells, we used flow cytometric analysis to measure propidium iodide fluorescence over the course of PaMx41Δ*acb2* phage infection. Propidium iodide (PI) is a membrane impermeable dye that enters cells with a porous or damaged membrane, binds to DNA and becomes fluorescent, thereby selectively labeling dead or dying cells. In contrast, intact cells do not uptake PI dye and metabolically active cells with minor membrane damage may uptake and pump out the dye and retain low levels of PI/red fluorescence. In Pa011 strains with endogenous or constitutive CBASS activity, <6% of PI/red fluorescence was detected over the course of phage infection compared to heat-killed control cells where ∼100% of cells effectively uptake and retain the PI dye (Figure 2D-E). In the Pa011 ΔCBASS strain, the cells slowly lysed over the eight-hour infection period (Figure 2C), so the PI dye was not effectively taken up by this cell population, or the cells were entirely lysed and could not retain any PI dye. We therefore relied on the heat-killed Pa011 cells as a positive control for PI-stained dead cells. These results support a model that Type II-A CBASS maintains cell viability while effectively stopping phage infection. Notably, we also observed that cells with increased 3’,3’-cGAMP production divide even more during phage infection compared to cells with endogenous 3’,’3’-cGAMP levels (Figure 2A, 2C), providing further evidence of a non-abortive outcome.

Due to our observations of a non-abortive outcome in bacteria with high, constitutive 3’,3’-cGAMP production, we hypothesized that CapV effector activity is antagonizing phage maturation, but not making the bacteria an inhospitable niche for phage replication *per se*. To test this, we used the PAO1 strain that naturally lacks CBASS because it is susceptible to a large diversity of phages. The full CBASS operon was integrated into the chromosome of PAO1 (PAO1^Pa011^) with its native promoter and additional copy of CdnA was supplied on a plasmid. We similarly observed robust cell growth during infection with CBASS sensitive phage JBD67Δ*acb2* (Figure S2B). When testing a large panel of phages using a solid infection assay, we observed that this strain with constitutive CBASS activity targeted at least eight genetically distinct phages, including some that encode the *acb2* inhibitor, reducing phage titer two to six orders of magnitude (Figure S3). These findings demonstrate that we can bypass both inhibitors and the need for phage activation of CBASS, revealing new phages that are sensitive to the activated CapV effector. However, certain phages, like PB-1, Ab06, and Ab09, remained resistant to CBASS across all conditions and highlight that CapV activity is phage-specific. These results can be further extended to the *Vibrio cholerae* CapV effector (Figure S3), which only shares 37.6% amino acid identity with *P. aeruginosa* CapV, but high structural similarity (Figure S3C). In a PAO1 strain expressing the *V. cholerae* DncV synthase and CapV effector, at least four genetically distinct phages were targeted three to five orders of magnitude (Figure S3), all of which were also targeted by *P. aeruginosa* CapV. Together, these data demonstrate that a CBASS effector acting on the cell membrane can be safely and constitutively activated in a manner that does not compromise cell viability and possesses specificity towards the targeted phages.

### Extracellular cyclic nucleotides induce CBASS-dependent cell growth

Given the apparent safety and efficacy of high, constitutive intracellular 3’3’-cGAMP levels, we hypothesized that this same effect could be achieved through supplementing media with exogenous signals. 3’,3’-cGAMP was titrated into the media and then Pa011 cells were infected with PaMx41Δ*acb2* phage (Figure 3A). Addition of ≤15 µM extracellular 3’,3’-cGAMP did not change cell growth during phage infection (Figure 3B). However, addition of ∼30 to 150 µM of extracellular 3’,3’-cGAMP rescued cell growth in the catalytically dead CdnA and Cap2 strains, which do not produce the signal endogenously, during phage infection (Figure S4A). Addition of any concentration of extracellular 3’,3’-cGAMP to the Pa011 strains without the CBASS operon, or with a catalytically dead CapV effector, did not improve cell growth during phage infection (Figure 3B, S4A), indicating that extracellular 3’,3’-cGAMP specifically improves CBASS anti-phage activity in a CapV-dependent manner. For context, 150 µM of extracellular 3’,3’-cGAMP is 10 to 100-fold more molecules than the intracellular 3’,3’-cGAMP detected in cells with CdnA overexpression. Under physiological conditions, we do not expect to detect this concentration of 3’,3’-cGAMP, but these results again demonstrate how safe and effective 3’,3’-cGAMP is for cells. Furthermore, adding extracellular 3’,3’-cGAMP to uninfected cells did not result in any cellular toxicity, but instead it modestly improved cell growth in a CapV-independent manner. Given that CBASS was previously shown to induce cell death within heterologous expression systems, importing cyclic nucleotide signals into uninfected cells would have been anticipated as a disadvantage to host survival. However, with our observation that CBASS activity does not compromise cell viability, the import of cyclic nucleotide signals would be a fitness advantage to the host and therefore warrants further investigation.

**Figure 3.**
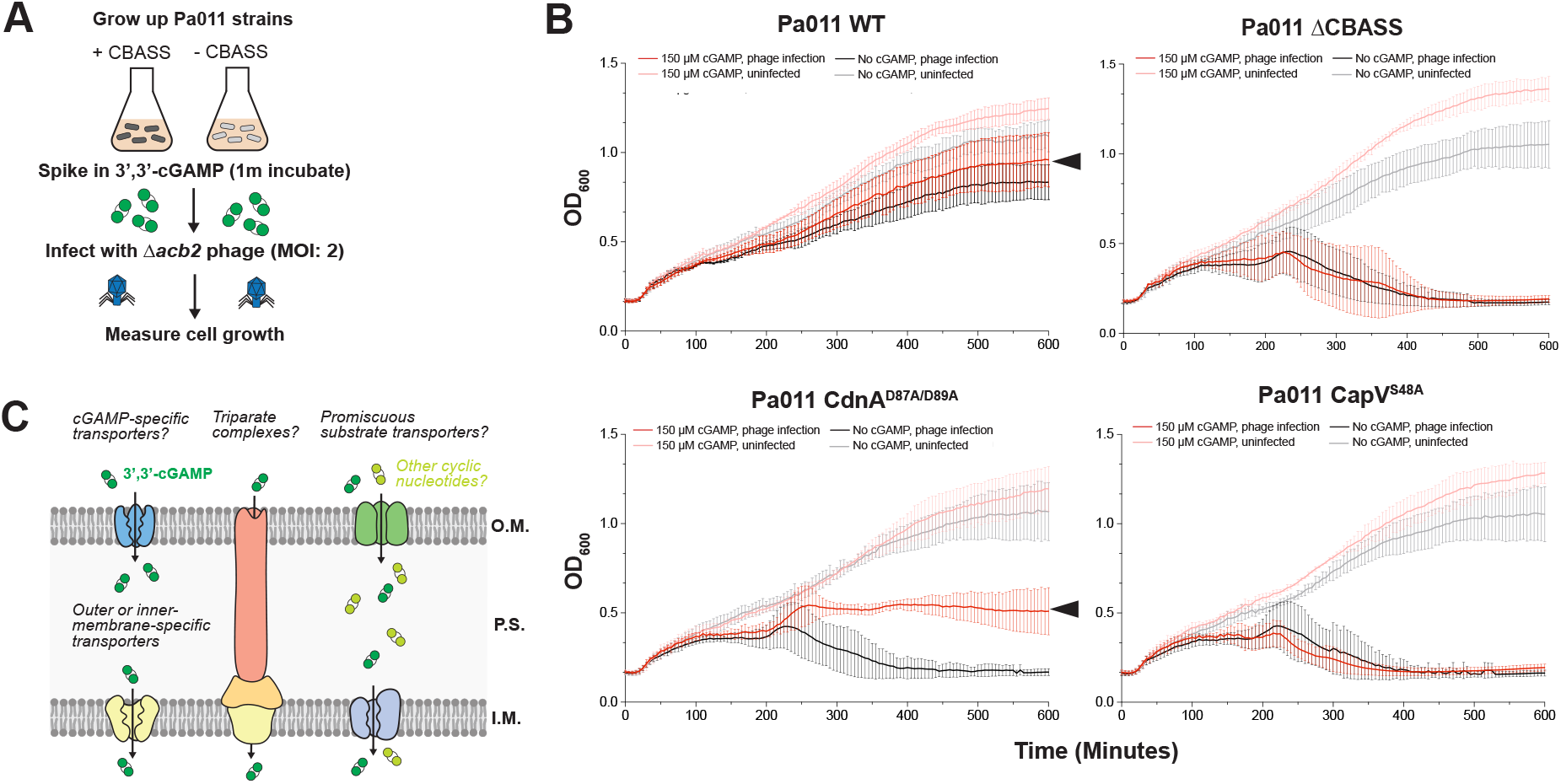
Extracellular 3’,3’-cGAMP signals rescue CBASS-dependent cell growth. **(A)** Preparation of bacterial cells. **(B)** OD_600_ measurements of Pa011 WT Type II-A CBASS, CdnA^D87A/D89A^ (dCdnA), and CapV^S48A^ (dCapV) strains infected with or without PaMx41Δ*acb2* phage (MOI:2) and with or without 3’,3’-cGAMP (∼150 µM) over 600 minutes (n=3 +/-s.d). **(C)** Model of putative cyclic nucleotide transporters in gram-negative bacteria.

### Type II-A CBASS limits phage DNA levels and mature virion production

To determine how CBASS limits phage replication without inducing cell death, we examined each stage of the phage replication cycle. Here, we used the Pa011 strains with endogenous CBASS expression for the molecular measurements. To start, we performed a single-step growth curve of PaMx41Δ*acb2* phage to evaluate the timing of the replication cycle and quantify mature virions. We observed that it takes 75-90 minutes from phage absorption to the detection of phage particles after chloroform treatment, which lyses the cells and ensures all mature virions are counted (Figure 4A). At the final time point (160 minutes post-infection), we observed that the Pa011 strain with endogenous CBASS activity (Pa011 WT) reduced the production of mature virions by ∼10-fold compared to cells lacking CBASS (Figure 4A). These data align with our findings presented earlier in this study (Figure 2B).

**Figure 4.**
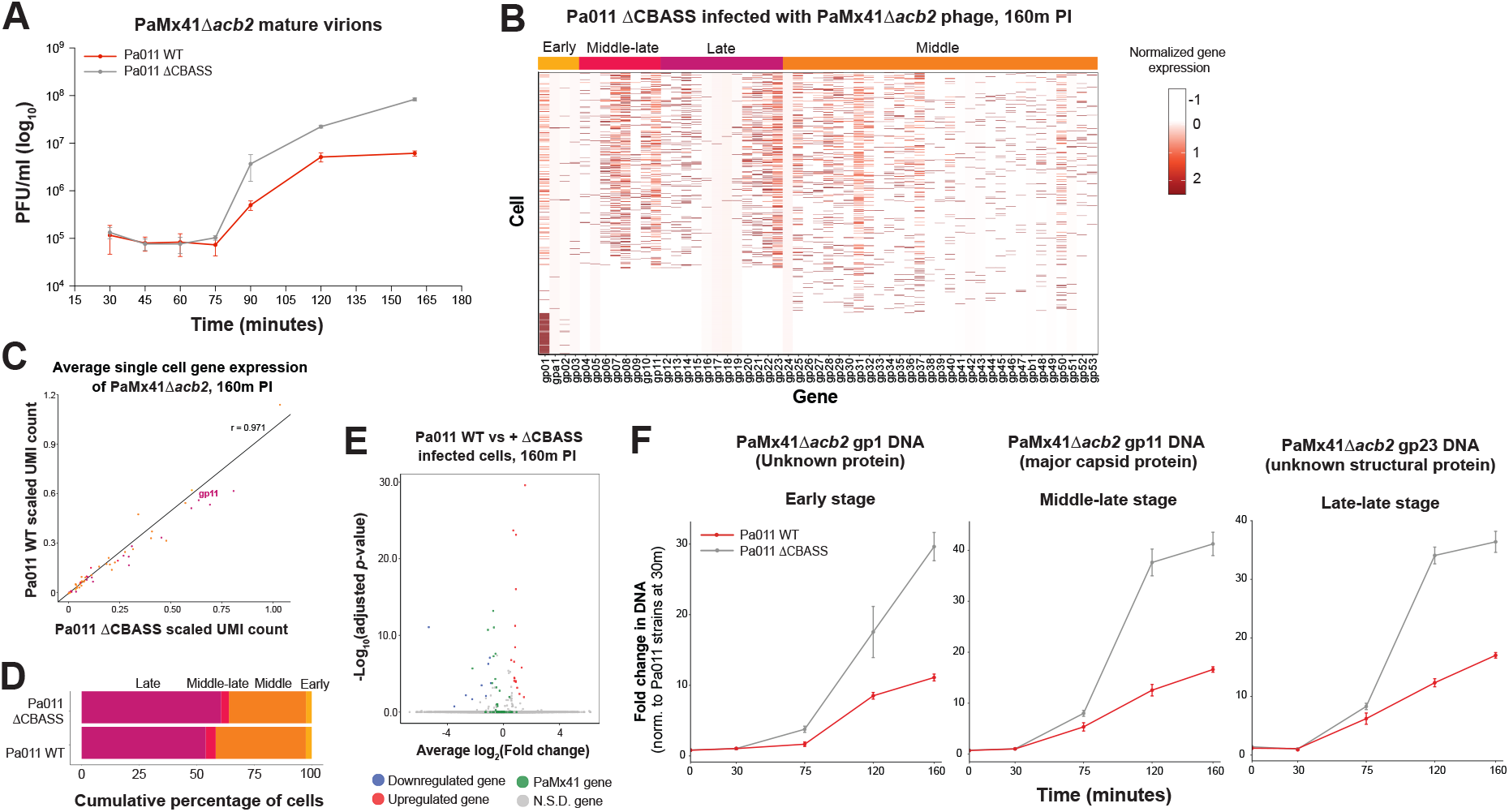
Type II-A CBASS limits PaMxΔ*acb2* phage DNA and mature virions. **(A)** Single-step growth curve of PaMx41Δ*acb2* phage infecting Pa011 WT or ΔCBASS strains (n=3 +/-s.d.). PFU/ml were quantified at the indicated times after chloroform treatment. **(B)** Single-cell RNA-Sequencing of the Pa011 ΔCBASS strain infected with PaMx41Δ*acb2* phage (MOI:2) at 160 minutes post-infection (PI). Phage genes were classified as early, middle, middle-late, and late based on Cruz-Plancarte et al. 2016. Note that gp24 represents acb2 and is not detectable. **(C)** Average single-cell phage gene expression levels in Pa011 WT verses Pa011 ΔCBASS strains infected with PaMx41Δ*acb2* (MOI:2, 160m PI). Gene expression in each cell scaled to the number of unique molecular identifiers (UMI). **(D)** Percentages of cells at each stage of phage infection in Pa011 WT and Pa011 ΔCBASS strains (MOI:2, 160m PI). **(E)** Pseudo-bulk RNA-Seq comparison of Pa011 WT verses Pa011 ΔCBASS strains infected with PaMx41Δ*acb2* (MOI:2, 160m PI). **(F)** DNA levels of PaMx41Δ*acb2* phage gp1 (early-stage gene, unknown protein), gp11 (middle-late stage; major capsid protein), and gp23 (late-late stage, unknown structural protein) during infection of Pa011 WT or ΔCBASS strains. Values were normalized to each respective Pa011 strains at 30 minutes PI.

To determine the stage that CBASS-sensitive phages fail to proceed through replication, we initially performed single-cell RNA-Sequencing (M3-seq)^29^ on Pa011 WT cells infected with PaMx41Δ*acb2* phage and measured early, middle, and late-stage gene expression. In the absence of CBASS, we observed that nearly all early, middle, and late gene expression states are detectable at 160 minutes post-infection (Figure 4B). This endpoint was chosen to maximize RNA detection. Next, we compared PaMx41Δ*acb2* phage gene expression in the presence and absence of endogenous CBASS activity. We found that phage gene expression correlated well between Pa011 WT and ΔCBASS strains (*r* = 0.971, Figure 4C), and there was no observable difference in the percentage of Pa011 cells at each stage of the phage replication cycle (Figure 4D). Pseudo-bulk RNA measurements also showed minimal or no significant changes in gene expression between Pa011 WT and ΔCBASS strains (Figure 4E). These collective results demonstrate that CBASS does not affect the phage transcriptome. This is consistent with previous work concluding that CBASS functions later in the phage replication cycle^10^.

Next, we hypothesized that CBASS may reduce phage virion production because there are fewer phage genomes overall. We measured phage DNA levels at three loci (gp1, gp11, and gp23) in the presence and absence of endogenous CBASS activity. DNA levels at each locus increased 30-40-fold in the Pa011 ΔCBASS strain, whereas DNA levels only increased ∼10-fold in the Pa011 strain, over 160 minutes post-infection (Figure 4F). This ∼3-fold decrease in maximum DNA levels is specific to CBASS-sensitive phages, such as PaMx41, because a genetically unrelated, CBASS-resistant phage Ab09 proceeds through its replication cycle across all Pa011 strains (Figure S5). Given the partial decrease of phage DNA levels, we suggest that phages proceed through DNA replication, but late-stage phage virion maturation and DNA packaging are impaired. As an alternative model to cell death, the CapV effector may interfere with procapsid assembly, which occurs at the cell membrane^30,31^, reducing virion production and DNA packaging.

## Discussion

To date, nearly all CBASS studies examining cyclic nucleotide-activated membrane effector proteins conclude that these effectors induce an abortive cell death mechanism^8,10,11,13,25^. The Type II-A CBASS system in *P. aeruginosa* strain BWHPSA011 (Pa011) features a phospholipase effector (CapV), which is activated by 3’,3’-cGAMP molecules and limits phage replication^16^. While this CBASS type can effectively limit the replication of sensitive phage, cell growth at the population-level is overwhelmed at high multiplicities of infection (MOIs). This observation complicates the ability to examine the cellular outcome of CBASS immunity because cells are dying from phage-induced lysis. However, we found that providing an extra copy of the CBASS synthase (CdnA) produces constitutive 3’,3’-cGAMP signals, which vastly enhances anti-phage activity and even overwhelms the CBASS inhibitor Acb2. In this case, constitutive 3’,3’-cGAMP signaling bypasses the need for phage activation. These findings align with previous studies with *Vibrio cholerae* and overexpressing the DncV synthase, or the full *V. cholerae* Type II-A CBASS operon, which produced ∼1.2 and 1.6 µM of intracellular 3’,3’-cGAMP^10,28^, respectively. In parallel, we observed that providing exogenous 3’,3’-cGAMP signals also activates CBASS, providing the first demonstration that importing cyclic nucleotide signals is safe and effective at protecting cells from phage infection. Future studies are needed to determine the mechanism of signal import.

A key result of this work is that a bacterial strain with constitutive CBASS activity maintains cell growth and viability at increasingly high MOIs (2-20) while blocking phage production. At the molecular level, we observed that the CBASS-sensitive phage, dsDNA lytic phage PaMx41, proceeded through most of its life cycle and exhibited partial reductions in DNA levels. We hypothesize that CapV activity at the cell membrane interferes with procapsid assembly, which limits virion biogenesis and leaves phage DNA unpackaged and vulnerable to degradation. This mechanism is motivated by previous studies that have shown that some phages assemble their procapsids at the inner membrane through an association between the phage portal, or portal adaptor proteins and the cell membrane, which serves as the anchor for the major capsid protein to begin assembling the procapsid structure^30,31^. Moreover, our previous study demonstrated that PaMx41 phage can escape CBASS targeting through mutations in its major capsid protein^16^, linking mutant capsid alleles to CBASS sensitivity, but the mechanism remains unknown.

While our study primarily relied on a single model phage, we also extended our work to a diverse panel of *P. aeruginosa* phages and it revealed the emergence of three phage classes: CBASS-sensitive, induced CBASS-sensitive, and CBASS-resistant. The first class of CBASS-sensitive phages are naturally targeted in the presence of CBASS expressed from the bacterial chromosome. The second class of CBASS-sensitive phages are targeted only upon CdnA overexpression. It is possible that these phages encode CBASS inhibitors, or weak/absent activators, but they are sensitive to CapV effector activity and become targeted with increased 3’,3’-cGAMP production. Notably, the *V. cholerae* CapV also restricted growth of a subset of these *P. aeruginosa* CapV-sensitive phages. The final class of phages remain resistant to CBASS even with enhanced 3’,3’-cGAMP levels. Altogether, these findings demonstrate that bacterial cells with activated CapV effectors are viable, and these cells also remain a hospitable niche for certain phages to proceed normally through their replication cycle. These phages may coordinate procapsid assembly at different locations in the cell that protect them from CapV-mediated modification of the cell membrane.

In summary, we have revealed that Type II-A CBASS, which is dependent on the function of the 3’,3’-cGAMP-activated CapV phospholipase effector, strikingly maintains cells growth and viability while stopping phage replication. Upon CapV activation, the bacterial cell limits replication of many, but not all, classes of phage, revealing a model of “effector specificity” that has been previously overlooked for cyclic nucleotide-activated CBASS effectors. Future work will examine how CapV-induced membrane perturbations interferes with procapsid assembly, which could more broadly apply to membrane-acting effectors across distinct bacterial immune systems.

## Acknowledgements

We thank Bondy-Denomy lab for their scientific insight and support for this project, including Dr. Wearn-Xin Yee for her expertise in bacterial cell viability assays. We thank the Gladstone Flow Cytometry Core for assisting with the flow cytometry experiments and access to their equipment. E.H. is supported by the National Institutes of Health (NIH) [5T32AI060537-20]. B.W. is supported by the NIH [T32HG003284] and the Charlotte Elizabeth Procter Fellowship from Princeton University. S.M. is supported by the National Institute of General Medical Sciences (NIGMS)–funded UCSF Medical Scientist Training Program (T32GM141323). J.B.-D. is supported by the Kleberg Foundation and previously by the NIH [R21AI168811], the Vallee Foundation, and the Searle Scholarship. Z.G. is supported by the NIH [1DP1AI190418 and 5R01GM143227].

## Author contributions

E.H. and J.B.-D. conceived the project and designed experiments. E.H. performed a majority of the *in vivo* bacterial cell experiments, with assistance from E.S. and S.M. who contributed to the cell growth and plaque assay experiments. B.W. performed all the phage DNA and RNA experiments, and bacterial host RNA experiments. E.H. wrote the rest of the manuscript and designed figures. All authors edited and provided feedback on the manuscript.

## Declaration of interests

J.B.-D. is a scientific advisory board member of SNIPR Biome and Excision Biotherapeutics, a consultant to LeapFrog Bio, and a scientific advisory board member and co-founder of Acrigen Biosciences and ePhective Therapeutics. Z.G. has a financial interest in ArrePath, Inc.

## Supplementary Figures

**Figure S1.**
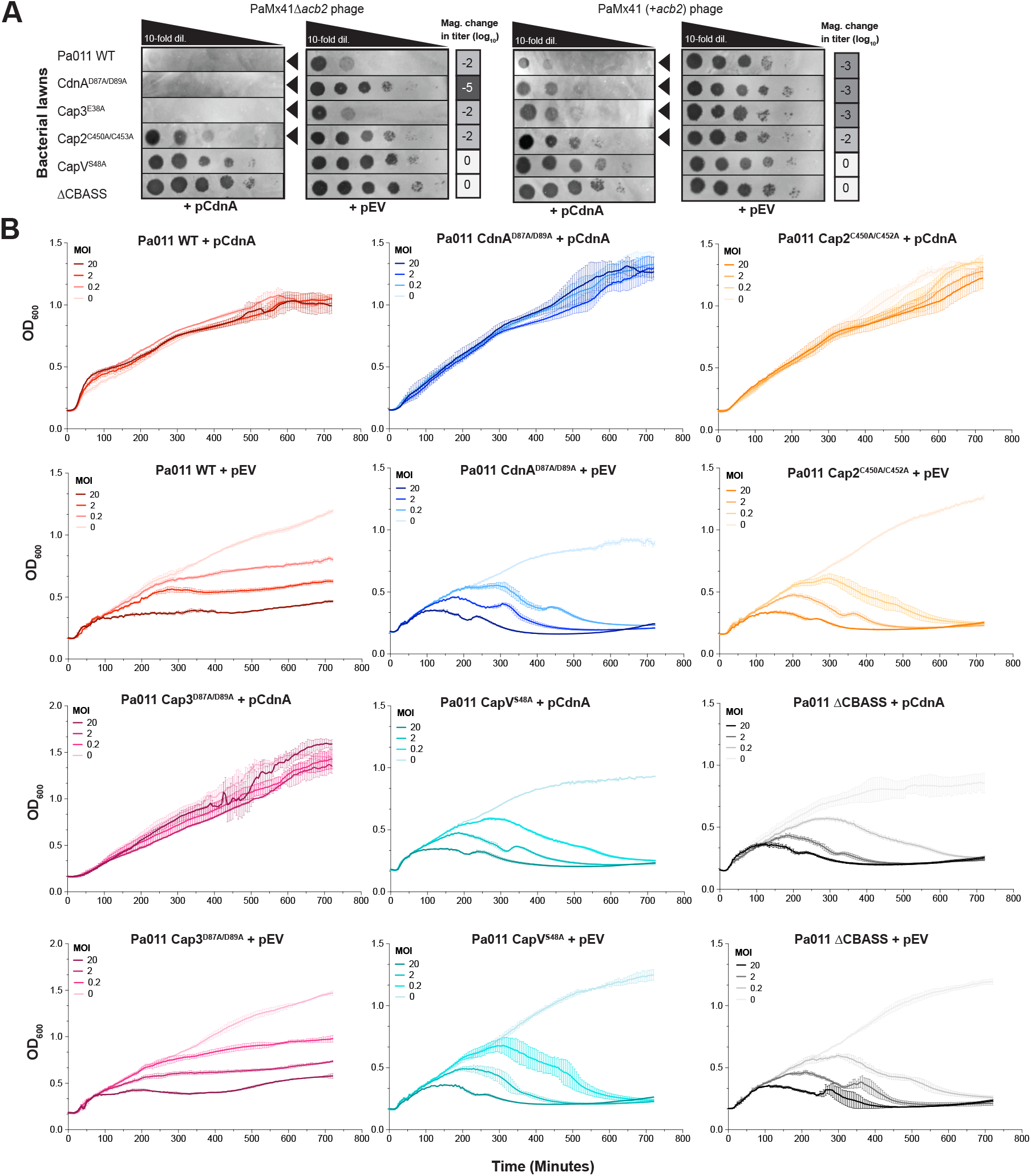
Increasing CdnA expression enhances anti-phage activity and cell growth. **(A)** PaMx41 phage that is CBASS sensitive (Δ*acb2*) or CBASS resistant (+*acb2*) phage spotted across lawns of Pa011 WT, ΔCBASS, or CBASS gene mutant cells expressing CdnA or empty vector (EV) plasmids. These plaque assays were used to quantify the order of magnitude change in phage titer by comparing the number of spots (with plaques, or clearings if plaques were not visible) on the CdnA overexpression strain divided by the EV strain (0.01% arabinose; n=3). Black arrowheads highlight notable reductions in phage titer. **(B)** OD_600_ measurements of Pa011 strains overexpressing CdnA or EV plasmids infected with an increasing multiplicity of infection (MOI) of PaMx41Δ*acb2* phage over 720 minutes (0.1% arabinose, n=1 biological replicate, n=3 technical replicates +/-s.d.).

**Figure S2.**
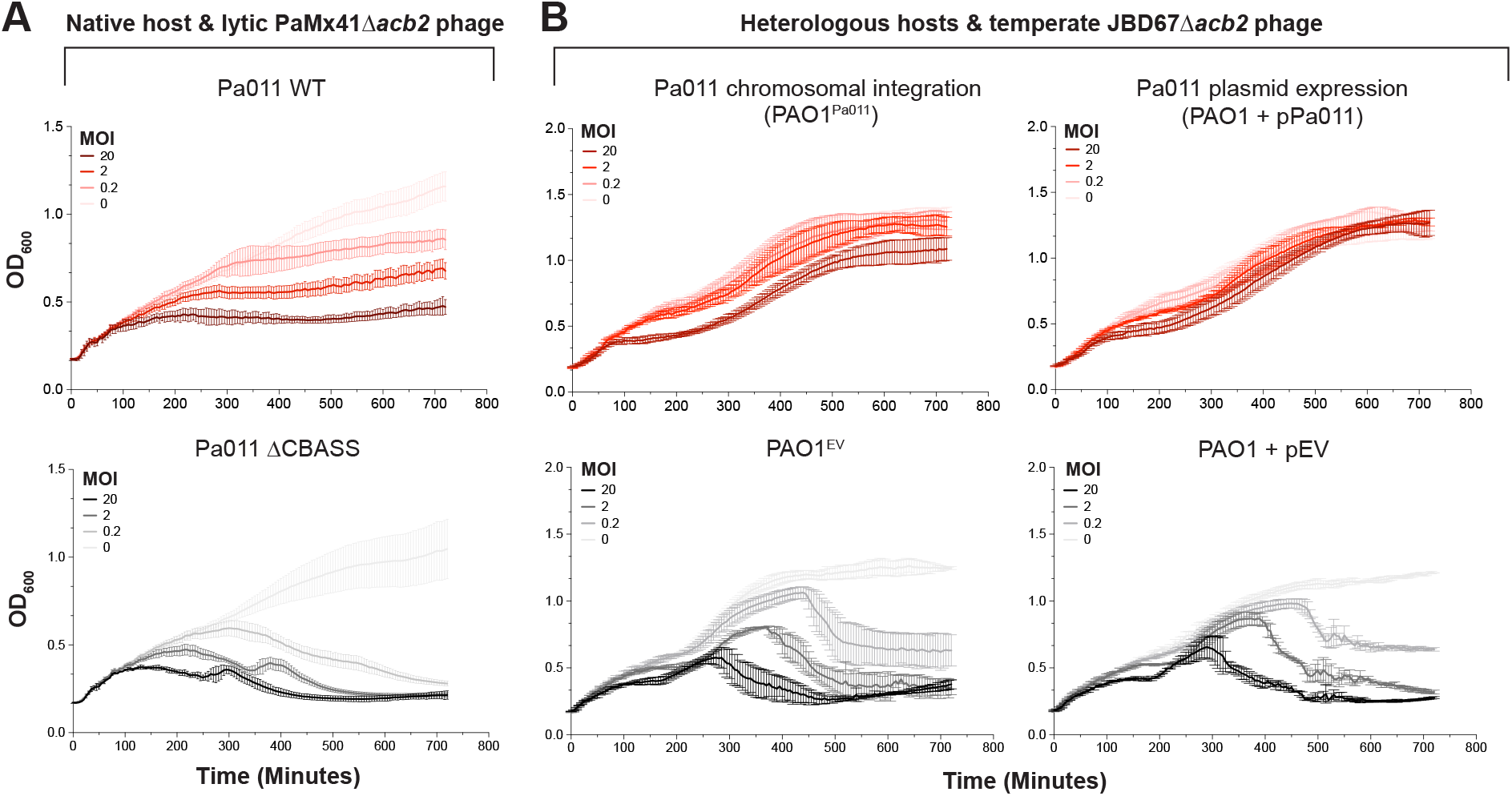
Heterologous expression of Type II-A CBASS enhances cell viability. OD_600_ measurements of **(A)** native Pa011 strains infected with an increasing multiplicity of infection (MOI) of lytic dsDNA PaMx41Δ*acb2* phage and **(B)** heterologous Pa011 strains infected with increasing MOIs of temperate dsDNA JBD67Δ*acb2* phage over 720 minutes (n=3 +/-s.d.). The Pa011 Type II-A CBASS operon was chromosomally integrated into the Tn7 site of the *Pseudomonas aeruginosa* strain PAO1 that naturally lacks CBASS (PAO1^Pa011^) and an empty Tn7 site (PAO1^EV^) served as the negative control (no IPTG induction needed). The Pa011 Type II-A CBASS operon was also expressed on a pHERD30T plasmid (PAO1 +pPa011) and empty vector (PAO1 + pEV) served as the negative control (0.1% arabinose induction.

**Figure S3.**
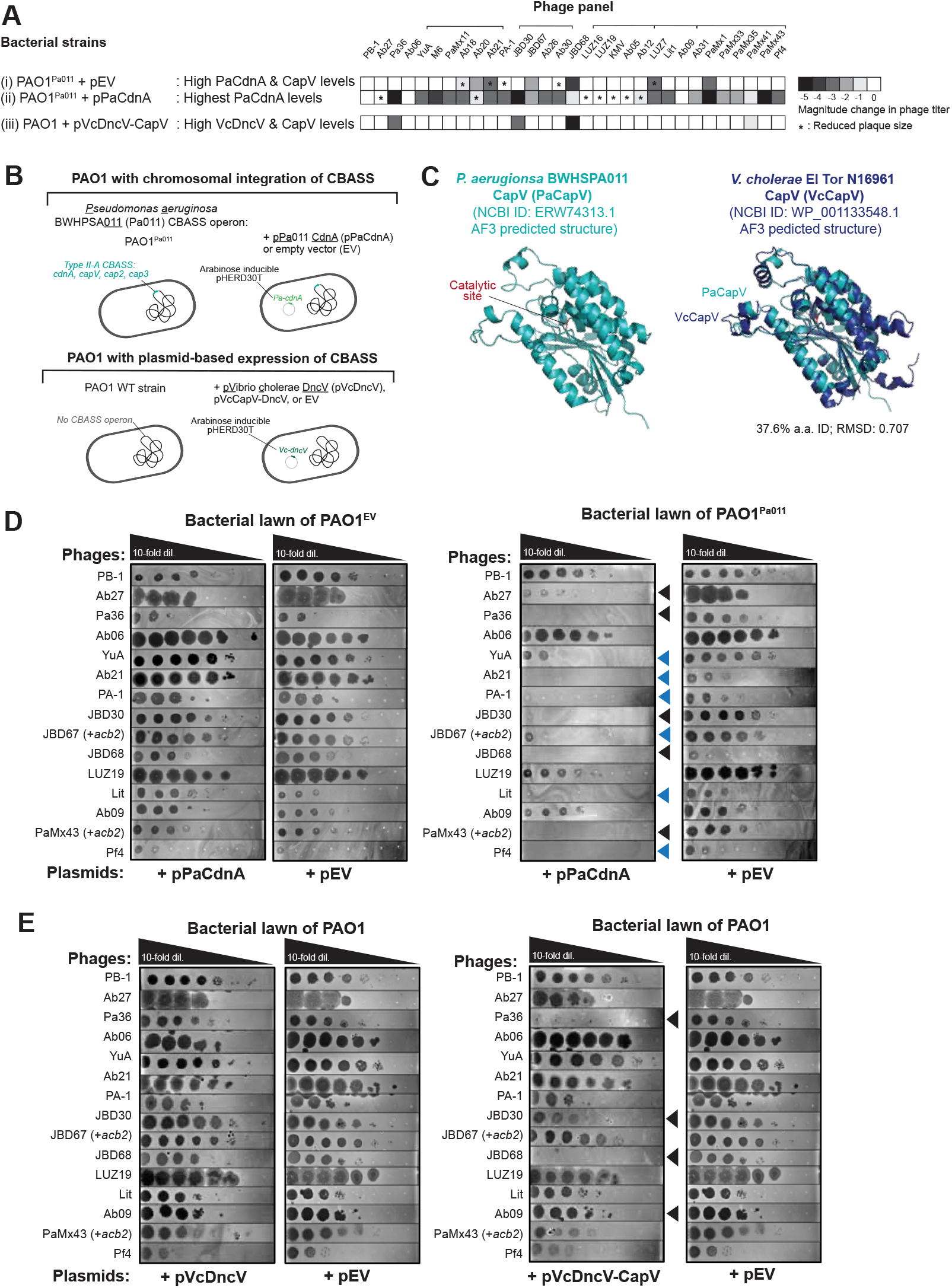
CapV effector activity exhibits phage specificity. **(A)** Matrix representing the order of magnitude change in phage titer on bacterial strains expressing full or partial Type II-A CBASS operons, and their relative expression levels of CBASS or synthase genes from *Pseudomonas aeruginosa* (Pa) or *Vibrio cholerae* (Vc). Titer changes were quantified by comparing the number of spots (with plaques, or clearings if plaques were not visible) between the following strains: (i) PAO1^Pa011^ + pEV strain (high PaCdnA and PaCapV expression levels) divided by PAO1^EV^ + pEV strain (no CBASS), (ii) PAO1^Pa011^ + pPaCdnA strain (highest PaCdnA expression levels) divided by PAO1^Pa011^ + pEV strain, and (iii) PAO1 + pVcDncV-CapV strain (high VcDncV and VcCapV expression levels) divided by PAO1 + pEV (no CBASS). Lines group phages with high genetic similarity. 68 total phages tested. Representative plaque assays of 15 genomically distinct phages shown. **(B)** Schematic of the strains used in these assays. **(C)** Structural comparison of CapV phospholipase effectors from *P. aeruginosa* BWHPSA011 (NCBI protein ID: ERW743131) or *V. cholerae* El Tor N16961 (NCBI: WP_001133548.1). Protein structures predicted using AlphaFold3. **(D-E)** Indicated phages spotted across lawns expressing CBASS synthase of empty vector (EV) plasmids (0.1% arabinose, n=1). Arrowheads highlight significant reductions in phage titer, and blue arrowheads indicated phages that are targeted by both *P. aeruginosa* and *V. cholerae* CBASS types.

**Figure S4.**
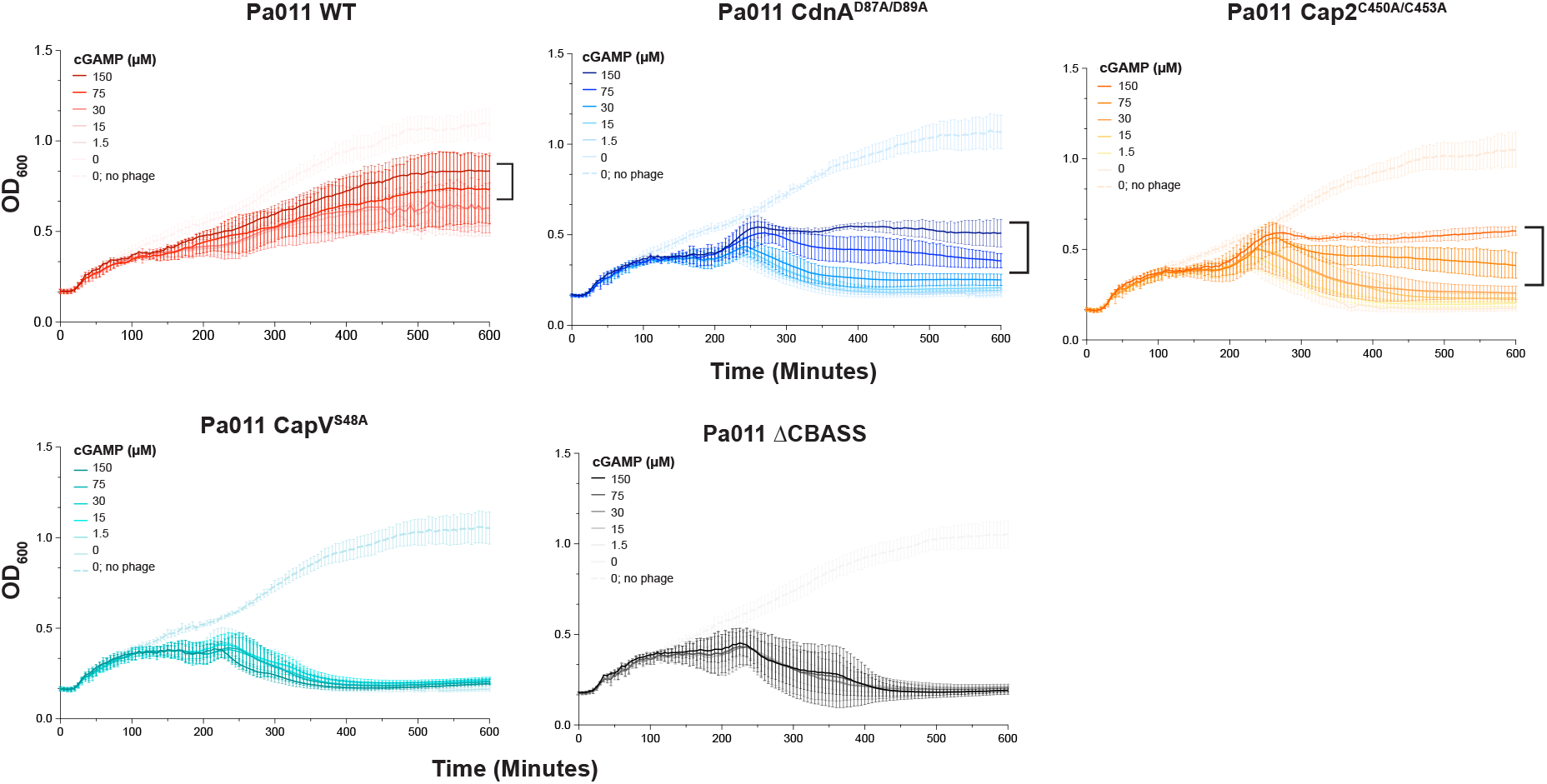
High levels of extracellular 3’,3’-cGAMP induces CBASS-dependent cell growth. OD_600_ measurements of native Pa011 WT, ΔCBASS, or CBASS mutant strains infected with or without PaMx41Δ*acb2* phage (MOI:2) and increasing concentrations of 3’,3’-cGAMP over 600 minutes (n=3 +/-s.d). Black brackets highlight increases in cell density relative to untreated cells.

**Figure S5.**
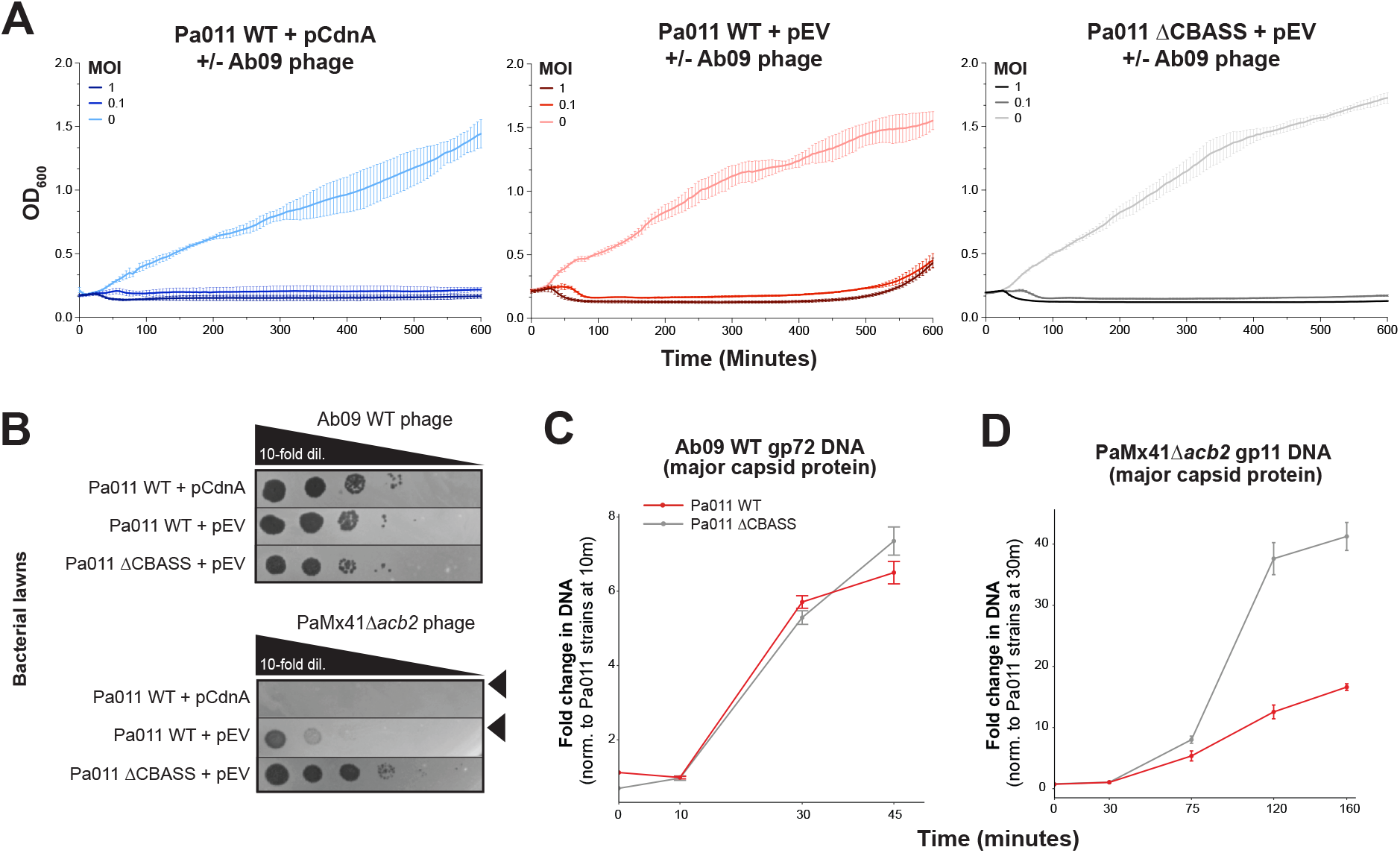
Reduction in DNA replication is specific to Type II-A CBASS-sensitive phage. **(A)** OD_600_ measurements of Pa011 strains expressing CdnA or empty vector (EV) plasmids infected with an increasing multiplicity of infection (MOI) of Ab09 WT phage over 600 minutes (0.1% arabinose, n=3 technical replicates +/-s.d.). **(B)** Ab09 WT and PaMx41Δ*acb2* phage spotted across lawns of Pa011 strains as described in (A) (0.1% arabinose, n=3 technical replicates +/-s.d.). **(C)** Ab09 WT phage gp72 (major capsid protein) DNA levels following infection of Pa011 WT or ΔCBASS strains. Values were normalized to each respective Pa011 strains at 10 minutes post-infection. **(D)** PaMx41Δ*acb2* phage gp11 (major capsid protein) DNA levels following infection of Pa011 strains Pa011 WT or ΔCBASS strains. Values were normalized to each respective Pa011 strains at 30 minutes post-infection.

**Figure S6.**
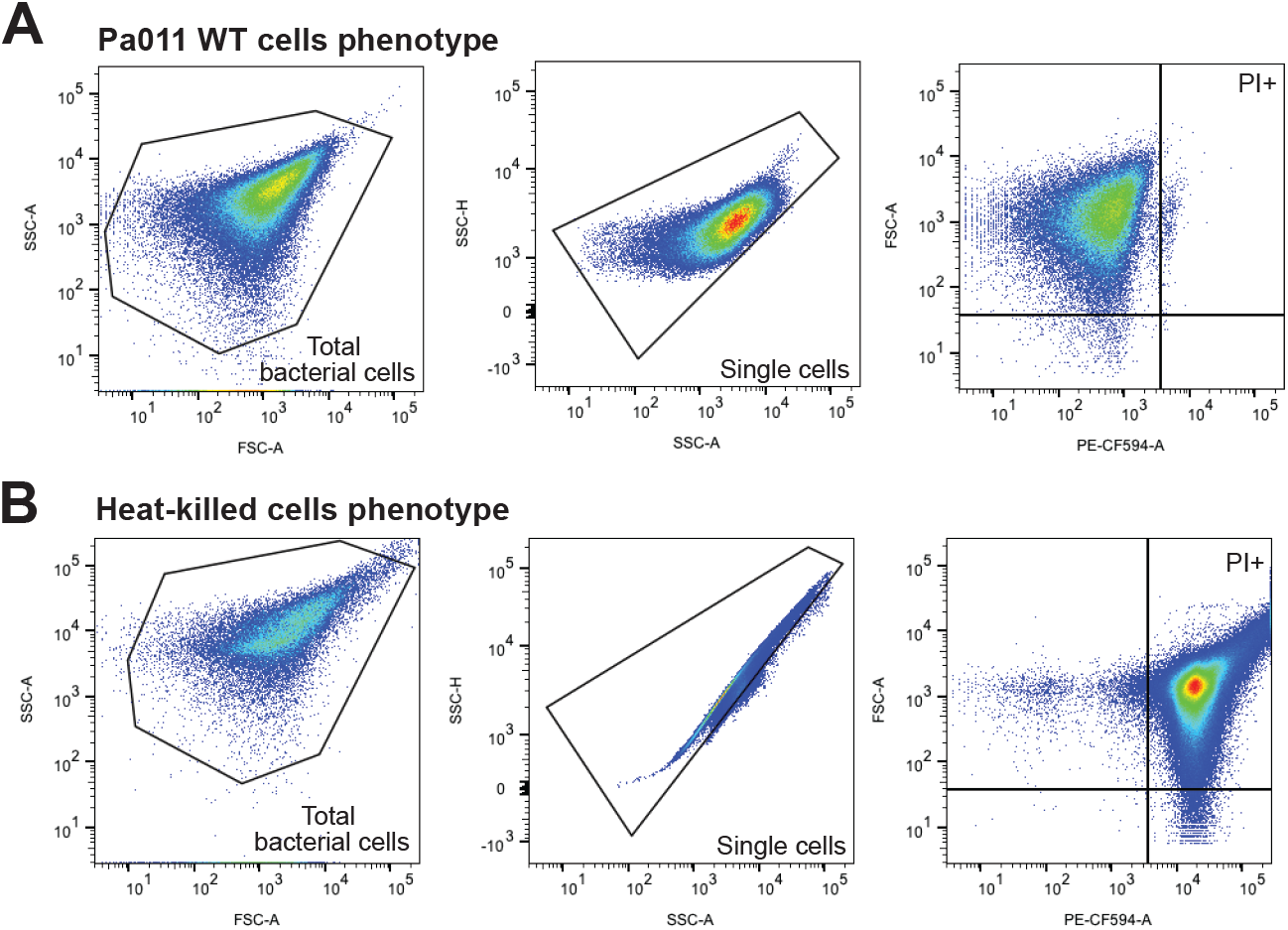
Flow cytometry gating strategy for propidium iodide expressing cells. **(A)** Gating strategy schematic for Propidium iodide (PI)-expressing bacterial cells and **(B)** heat-killed positive control (dead) cells. Note that cell shape (SSC-H v SSC-A) differs significantly between healthy (A) and dead (B) cells in *P. aeruginosa*. This was observed with over three biological replicates.

## Resources availability

### Lead Contact

Further information and requests for resources and reagents should be directed to and will be fulfilled by the lead contact, Joseph Bondy-Denomy.

### Materials Availability

All unique/stable reagents generated in this study are available from the Lead Contact with a completed Materials Transfer Agreement.

### Experimental model and subject details

#### Bacterial strains and phages

The *P. aeruginosa* strains (BWHPSA011 and PAO1) and *E. coli* strains (DH5ɑ and SM10) were grown in Lysogeny broth (LB) medium at 37°C both with aeration at 175 r.p.m. Plating was performed on LB solid agar with 10 mM MgSO_4_ when performing phage infections, and when indicated, gentamicin (50 µg/ml for *P. aeruginosa* and 15 µg/ml for *E. coli*) was used to maintain the pHERD30T plasmid. Gene expression of pHERD30T constructs was induced by the addition of L-arabinose, and gene expression of chromosomal insertions was induced by isopropyl-β-D-thiogalactopyranoside (IPTG). Final concentrations are indicated in methods and figure legends.

## Method details

### CBASS operons

The *P. aeruginosa* BWHPSA011 (Pa011) strain (NCBI Reference Sequence ID: NZ_AXQR00000000.1) contains a Type II-A CBASS operon, with CapV (NCBI Nucleotide ID: Q024_03602, Protein ID: P0DX85.1), CdnA (NCBI Nucleotide ID: Q024_03601, Protein ID: P0DX86.1), Cap2 (NCBI Nucleotide ID: Q024_03600, Protein ID: P0DX87.1), and Cap3 (No NCBI ID, intergenic region, protein sequence: MQFMSSWAADDNRTLLHFSKSTLETFRQHVQASDSDC EAGGLLLGSVHGTHMLIEHATVPTAWDKRFRYLFERMPFGHEAIALARWTASQGTIRHLGEWHTHPEDNPNPSGLDRSEWNRLSAKRRDKRPTLAVIVGRNALYIELVPGSGQGSVFSPVE).

### Recombinant DNA in bacterial cells

The shuttle vector that replicates in *P. aeruginosa* and *E. coli*, pHERD30T^32^ was used for cloning and episomal expression of genes in *P. aeruginosa* BWHPSA011 (Pa011) or ATCC 27853 (Pa278). Pa011 CdnA, Pa011 Type II-A CBASS operon (CapV, CdnA, Cap3, Cap2) described above were cloned with ∼50bp upstream. The PaMx41 capsid constructs were cloned directly downstream of the promoter. A final arabinose concentration of 0.05% or 0.1% was used to induce expression of each construct. This vector has an arabinose-inducible promoter and a selectable gentamicin marker. Vector was digested with SacI and PstI restriction enzymes unless stated otherwise and then purified. Inserts were amplified by PCR using bacterial overnight culture or phage lysate as the DNA template, and joined into the pHERD30T vector at the SacI-PstI restriction enzyme cut sites by Hi-Fi DNA Gibson Assembly (NEB) following the manufacturer’s protocol. The resulting plasmids were transformed into *E. coli* DH5ɑ. All plasmid constructs were verified by sequencing using primers that annealed to sites outside the multiple cloning site. *P. aeruginosa* cells were transformed with the pHERD30T constructs using electroporation or conjugation via SM10 *E. coli* cells.

### Chromosomal CBASS integration

For chromosomal insertion of the Pa011 CBASS operon, the integrating vector pUC18-mini-Tn7T-LAC^33^ and the transposase expressing helper plasmid pTNS3^34^ were used to insert the Pa011 Type II-A CBASS operon at the Tn7 locus in *P. aeruginosa* PAO1 strain (PAO1^Pa011^). The Pa011 operon was cloned with ∼250bp upstream and 0 mM IPTG and 0.3 mM IPTG was necessary to induce expression of each respective operon. The pUC18-mini-Tn7T-LAC empty vector (E.V.) served as the control strain (Pa^EV^). The vector was linearized using around-the-world PCR, treated with DpnI, and then purified. Two overlapping inserts encompassing the CBASS operon were amplified by PCR using Pa011 overnight culture as the DNA template, and joined into the pUC18-mini-Tn7T-LAC vector a the SacI-PstI restriction enzyme cut sites by Hi-Fi DNA Gibson Assembly (NEB) following the manufacturer’s protocol. The resulting plasmids were used to transform *E. coli* DH5ɑ. All plasmid constructs were verified by sequencing using primers that annealed to sites outside the multiple cloning site. *P. aeruginosa* PAO1 cells were electroporated with pUC18-mini-Tn7T-LAC and pTNS3 and selected for on gentamicin. Potential integrants were screened by colony PCR with primers PTn7R and PglmS-down, and then verified by sequencing using primers that anneal to sites outside the attTn7 site. Electrocompetent cell preparations, transformations, integrations, selections, plasmid curing, and FLP-recombinase-mediated marker excision with pFLP were performed as described previously^33^.

### Phage growth

All phages were grown at 37°C with solid LB agar plates containing 20 ml of bottom agar containing 10 mM MgSO_4_ and antibiotics or inducers, as appropriate. Phages were grown on a permissible *P. aeruginosa* host, such as Pa011 ΔCBASS or PAO1 WT, which lack CBASS. Whole plate infections were performed with 150 µl of overnight cultures infected with 10 µl of low titer phage lysate (>10^4-7^ PFU/ml), and then mixed with 3 ml of 0.7% top agar 10 mM MgSO_4_ for plating on the LB solid agar. After incubating 37°C overnight, SM buffer was added until the solid agar lawn was completely covered and then incubated for 5-10 minutes at room temperature. The whole cell lysate was collected, and a 1% volume of chloroform was added, and then left to shake gently on an orbital shaker at room temperature for 15 min followed by centrifugation at maximum g for 1 min to remove cell debris. The supernatant phage lysate was stored at 4°C for downstream assays.

### Plaque assays

Plaque assays were conducted at 37°C with solid LB agar plates. 150 µl of overnight bacterial culture was mixed with top agar and plated. Phage lysates were diluted 10-fold in SM buffer and then 1.5 µl spots were applied to the top agar after it had been poured and solidified.

### Intracellular 3’,3’-cGAMP measurements of bacterial cells

Cell lysates were prepared similarly to previous methods^10,16^, in which *P. aeruginosa* BWHPA011 (Pa011) cells harboring a catalytically dead *capV* gene (CapV^S48A^) were used and then transformed with a pHERD30T vector expressing Pa011 CdnA or an empty vector (EV) control. Cells were taken from overnight culture, diluted 1:100 in 150ml LB medium with 10 mM MgSO4 added, Gentamycin (50 µg/ml), and Arabinose (0.1%) in a 500ml flask, and then grown at 37°C (175 r.p.m.) until reaching an OD_600nm_ of ∼0.3. From the culture, 50ml of cells were kept uninfected and 50ml of cells were then infected with PaMx41Δ*acb2* to obtain an MOI of ∼5 and ensure at least one or more phages were infecting each bacterial cell. After 60 minutes following infection, the culture was centrifuged at 7,500 g for 10 mins at 4°C. Following centrifugation, supernatant was removed and pellets were kept on ice until resuspended in 600 µl of phosphate buffer (50 mM sodium phosphate (pH 7.4), 300 mM NaCl, 10% (v/v) glycerol). The resuspended pellet was supplemented with 1 µl hen-lysozyme (Sigma-Aldrich), vortexed briefly, and incubated at 25°C for 10 min. The resuspended cells were then mixed with Lysing Matrix B (MP) beads and cells were disrupted mechanically using Mini-Beadbeater 16 Biospec Products (1 cycle of 2:30, 3,450 oscillations/m, at 4 °C). Cell lysates were then centrifuged at 16,000 g for 10 min at 4°C. For each condition, the supernatant was loaded onto a 3kDa filter (Amicon Ultra-0.5 centrifugal filter unit; Merk) and then centrifuged at 16,000 g for 45 min at 4 °C. The flow-through (containing small molecules less than 3kDa) was used as the sample for 3’,3’-cGAMP measurements, and each sample was run in technical triplicate on a 3’,3’-cGAMP ELISA Kit (Arbor Assays) and standards were prepared in the same phosphate buffer. 3’,3’-cGAMP concentrations were calculated using a sigmoidal standard curve via GraphPad Prism (v 9.4.1).

### OD_600_ cell growth curve experiments

Overnight cultures were diluted 1:100 in 25 ml LB medium with 10 mM MgSO4, antibiotics and inducers, as appropriate (150ml flask), and then grown at 37°C (175 r.p.m.) until reaching an OD_600_ of ∼0.3. From the culture, 140 µl was aliquoted in a 96-well plate and infected with <10 µl of phage to reach a final MOI of 20, 2, and 0.2. Each infection was performed in technical triplicate, and cultured with maximum double orbital rotation at 37 °C for 12h with OD_600_ measurements taken every 5 minutes using a BioTek Synergy H1 microplate reader (Gen5 software).

### CFU and PFU measurements

Overnight cultures were diluted 1:100 in 25 ml LB medium with 10 mM MgSO4, antibiotics and inducers, as appropriate (150ml flask), and then grown at 37 °C (175 r.p.m.) until reaching an OD_600_ of ∼0.3. From the culture, 3 ml aliquots were prepared in 6 ml glass culture tubes. One aliquot was kept uninfected and another infected with < 20 µl of phage to reach a final MOI of 2. Cultures were incubated at 37 °C (90 r.p.m.) using a Fisherbrand Isotemp shaking water bath. Pre-infection (0h) and then 2-, 4-, 6-, and 8-hours post-infection aliquots were taken to measure colony forming units (CFU), plaque forming units (PFU), and single-cell flow cytometry measurements (described below). For CFU counts, 1 or 10 µl of culture was diluted in SM buffer to prevent further growth and then plated on solid agar with antibiotics and inducers, as appropriate. For PFU counts, 50 µl of culture was collected with 5 µl of chloroform, centrifuged at maximum g for 1 min, and then stored at 4°C. Whole plate infections were performed as described above. After incubating 37°C overnight, individual colonies or plaques were counted and CFU/ml or PFU/ml calculated, respectively.

### Single-step growth curve experiments

Overnight cultures were diluted 1:100 in 25 ml LB medium with 10 mM MgSO4 (150ml flask) and supplemented with 0.05% arabinose, and then grown at 37°C (175 r.p.m.) until reaching an OD_600_ of ∼0.3. From the culture, 2 ml aliquots of cells were centrifuged at 9,000 g for 3 minutes and supernatant removed. Cells were resuspended in 200 µl LB medium with 10 mM MgSO4. One aliquot was kept uninfected and another infected with phage to reach a final MOI of 0.2. Cells were kept on ice for 20 minutes to effectively absorb phage. Cells were resuspended with medium to reach 1ml and then centrifuged at 9,000 g for 3 minutes. Supernatant carefully removed to eliminate unabsorbed phage, and resuspended in 1 ml of fresh LB medium with 10 mM MgSO4. An aliquot of phage was taken here as the first time point (T:0m). The remaining culture was transferred to a 3 ml glass culture tube and incubated at 37°C (90 r.p.m.) using a Fisherbrand Isotemp shaking water bath. At 0, 20-, 40-, 60-, 80-, and 120-minutes post-infection 50 µl aliquots were taken and added to 5 µl of chloroform, spun down at maximum g for 1 min, and then stored at 4°C. Whole plate infections were performed as described above. After incubating 37°C overnight, individual colonies or plaques were counted and CFU/ml or PFU/ml calculated, respectively. Frequency of surviving phage was calculated using the PFU/ml of output phage normalized to PFU/ml input phage.

### Single-cell RNA sequencing

Overnight cultures were diluted 1:100 in LB medium with 10 mM MgSO4 and grown to an OD of ∼0.3. 4mL aliquots of the culture were taken, and PaMx41 variants were added to an MOI of 2. Aliquots of 1mL of culture were taken 60-, 160-, and 360-minutes post infection. For each aliquot, cells were spun down for 2 minutes at 5,000g at room temperature. Cells were then fixed by resuspending in 4mL of 4% formaldehyde and left rotating overnight. Cells were then prepared for single-cell sequencing following M3-Seq^29^. We loaded 3 lanes of the Chromium Next GEM Chip H (10x Genomics, 1000162) with 280,000 cells in each lane. In-droplet ligation, and subsequent library preparation was performed following the M3-Seq protocol, in which we used the Pseudomonas rRNA probes to deplete Pa011 rRNA. Libraries were then pooled and sequenced on the NovaSeq S2 100 cycle kit (Illumina 20028316) with the following read structure: 26bp Read 1, 30bp i5 index, 8bp i7 index, 74bp Read2. Following sequencing and demultiplexing, reads were mapped to a combined Pa011 and PaMx41 genome. Demultiplexing, cell-calling, and analyses were performed following the protocols described in the M3-Seq paper. Briefly, we used Seurat (v.5.0.3)^35^ for all custom downstream analyses. For the phage gene expression analysis, we created matrices containing only phage genes and filtered the matrix to have cells that had more than or equal to one phage unique molecular identifiers (UMI). To identify which stage of phage infection each cell was in, we first classified all the PaMx41 genes as early, middle, middle-late, or late following northern blot analyses in the original paper characterizing PaMx41^36^. We then iteratively assigned cells to each stage by identifying the “latest” phage gene that was expressed in each cell.

### qPCR, RT-qPCR, and Bulk RNA-Seq

Overnight cultures were diluted 1:100 in LB medium with 10 mM MgSO4 and supplemented with 0.05% arabinose, and then grown at 37°C (175 r.p.m.) until reaching an OD_600_ of ∼0.3. From the culture, 4 ml aliquots of cells were infected with phage to reach a final MOI of 2. An aliquot of 1 ml was immediately taken (T:0m) and then up to 160-minutes post-infection. Cells were pelleted at maximum g for 90 sec, supernatant removed, and then lysozyme added to 15mg/mL. From these pellets, total DNA was extracted using Qiagen DNeasy Blood & Tissue Kit (Cat. No. 69504), and total RNA was extracted using SPRI-select beads, following the RNAdvance Cell V2 protocol (Cat. No. A47943). RNA for the 160-minute timepoints were sent to SeqCenter for library prep and RNA-Sequencing (12M reads, paired end with rRNA depletion). For RT-qPCR, RNA was reverse transcribed using NEB’s LunaScript RT SuperMix (Cat No. E3010). We then used the ThermoFisher PowerUp SYBR Green (Cat No. A25742) for qPCR on the reverse transcribed cDNA and the extracted total DNA. Both reactions were performed on the ViiA 7 thermocycler. Primers targeting the *Pseudomonas aeruginosa* 5s ribosomal RNA gene was used the internal control during calculations, and gene fold changes were normalized against the readout of the Pa011 ΔCBASS strain at 30m post-infection.

### Flow cytometry

Cultures were prepared as described above. Pre-infection, 1 ml of culture was collected and served as the first time point (T:0h) and another 1 ml served as the heat-killed positive control (95°C for 10 min). At 2-, 4-, 6-, and 8-hours post-infection, 200 µl of culture was collected. For each sample, an equal volume of diluted Propidium Iodide (PI) was added (1:1000 in 1x PBS, Thermo Fisher, LIVE/DEAD Fixable Red Dead Cell Stain Kit, L34971), briefly vortexed, and then incubated for 30 min at room temperature, shaking, and covered from light. Cells were centrifuged at 9,000 g for 3 mins, supernatant removed, and resuspended in 200 µl of 3% PFA. Cells were briefly vortexed, incubated for 15 min at room temperature and covered from light. Cells were centrifuged at 9,000 g for 3 mins, supernatant removed, and resuspended in 1 ml 1x PBS (twice). Cells resuspended in 200 µl of PBS and stored overnight at 4°C and covered from light. Cells were then analyzed on a BD LSRFortessa X-20 (BD Biosciences) using BD FACSDiva software version 9.0.0 with >20,000 post-gating single cell events recorded for each sample. PI+ cells was calculated using the frequency of events (cells) in the PI-expressing population out of the single-cell population. The data was analyzed with FlowJo (v10.10.0) and gating strategy is shown in Figure S6.

### Extracellular 3’,3’-cGAMP experiments

Cells were prepared and measured as described for the OD_600_ experiments. However, from the culture, 140 µl was aliquoted in a 96-well plate and then 5 to 15 µl of diluted 3’,3’-cGAMP (Sigma Aldrich, SML1232) was added to reach a final concentration of 1 to 100 µg/ml of cGAMP. Cells were incubated at room temperature for 1 minute. Afterwards <10 µl of phage to reach a final MOI of 2. Each infection was performed in biological replicate.

### Quantification and statistical analyses

Statistical details for each experiment can be found in the figure legends and outlined in the corresponding methods details section. Data are plotted with error bars representing standard deviation (s.d.) as indicated in the legends.

